# Modeling error in experimental assays using the bootstrap principle: Understanding discrepancies between assays using different dispensing technologies

**DOI:** 10.1101/033985

**Authors:** Sonya M. Hanson, Sean Ekins, John D. Chodera

## Abstract

All experimental assay data contains error, but the magnitude, type, and primary origin of this error is often not obvious. Here, we describe a simple set of assay modeling techniques based on the bootstrap principle that allow sources of error and bias to be simulated and propagated into assay results. We demonstrate how deceptively simple operations—such as the creation of a dilution series with a robotic liquid handler—can significantly amplify imprecision and even contribute substantially to bias. To illustrate these techniques, we review an example of how the choice of dispensing technology can impact assay measurements, and show how large contributions to discrepancies between assays can be easily understood and potentially corrected for. These simple modeling techniques—illustrated with an accompanying IPython notebook—can allow modelers to understand the expected error and bias in experimental datasets, and even help experimentalists design assays to more effectively reach accuracy and imprecision goals.

## I. INTRODUCTION

Measuring the activity and potency of compounds— whether in biophysical or cell-based assays—is an important tool in the understanding of biological processes. However, understanding assay data for the purpose of optimizing small molecules for use as chemical probes or potential therapeutics is complicated by the fact that all assay data are contaminated with error from numerous sources.

Often, the dominant contributions to assay error are simply not known. This is unsurprising, given the number and variety of potential contributing factors. Even for what might be considered a straightforward assay involving fluorescent measurements of a ligand binding to a protein target, this might include (but is by no means limited to): compound impurities and degradation [1–4], imprecise compound dispensing [5, 6], unmonitored water absorption by DMSO stocks [4], the effect of DMSO on protein stability [7], intrinsic compound fluorescence [8, 9], compound insolubility [10] or aggregation [9, 11–14], variability in protein concentration or quality, pipetting errors, and inherent noise in any fluorescence measurement—not to mention stray lab coat fibers as fluorescent contaminants [15]. Under ideal circumstances, control experiments would be performed to measure the magnitude of these effects, and data quality tests would either reject flawed data or ensure that all contributions to error have been carefully accounted for in producing an assessment of error and confidence for each assayed value. Multiple independent replicates of the experiment would ideally be performed to verify the true uncertainty and replicability of the assay data^1^.

Unfortunately, by the time the data reach the hands of a computational chemist (or other data consumer), the opportunity to perform these careful control experiments has usually long passed. In the worst case, the communicated assay data may not contain any estimate of error at all. Even when error has been estimated, it is often not based on a holistic picture of the assay, but may instead reflect historical estimates of error or statistics for a limited panel of control measurements. As a last resort, one can turn to large-scale analyses that examine the general reliability of datasets across many assay types [17, 18], but this is to be avoided unless absolutely necessary.

When multiple independent measurements are not available, but knowledge of how a particular assay was conducted *is* available, this knowledge can inform the construction of an assay-specific model incorporating some of the dominant contributions to error in a manner that can still be highly informative. Using the *bootstrap principle—*where we construct a simple computational replica of the real experiment and simulate virtual realizations of the experiment to understand the nature of the error in the experimental data—we often do a good job of accounting for dominant sources of error. Using only the assay protocol and basic specifications of the imprecision and inaccuracy of various operations such as weighing and volume transfers, we show how to construct and simulate a simple assay model that incorporates these important (often dominant) sources of error. This approach, while simple, provides a powerful tool to understand how assay error depends on both the assay protocol and the imprecision and inaccuracy of basic operations, as well as the true value of the quantity being measured (such as compound affinity). This strategy is not limited to computational chemists and consumers of assay data—it can also be used to help optimize assay formats before an experiment is performed, help troubleshoot problematic assays after the fact, or ensure that all major sources of error are accounted for by checking that variation among control measurements match expectations.

We illustrate these concepts by considering a recent example from the literature: a report by Ekins et al. [19] on how the choice of dispensing technology impacts the apparent biological activity of the same set of compounds under otherwise identical conditions. The datasets employed in the analyses [20, 21] were originally generated by AstraZeneca using either a standard liquid handler with fixed (washable) tips or an acoustic droplet dispensing device to prepare compounds at a variety of concentrations in the assay, resulting in highly discrepant assay results (Figure 1). The assay probed the effectiveness of a set of pyrimidine compounds as anti-cancer therapeutics, targeting the EphB4 receptor, thought to be a promising target for several cancer types [22, 23]. While the frustration for computational modelers was particularly great, since quantitative structure activity relationship (QSAR) models derived from these otherwise identical assays produce surprisingly divergent predictions, numerous practitioners from all corners of drug discovery expressed their frustration in ensuing blog posts and commentaries [24–26]. Hosts of potential explanations were speculated, including sticky compounds absorbed by tips [27] and compound aggregation [13, 14].

**FIG. 1.**
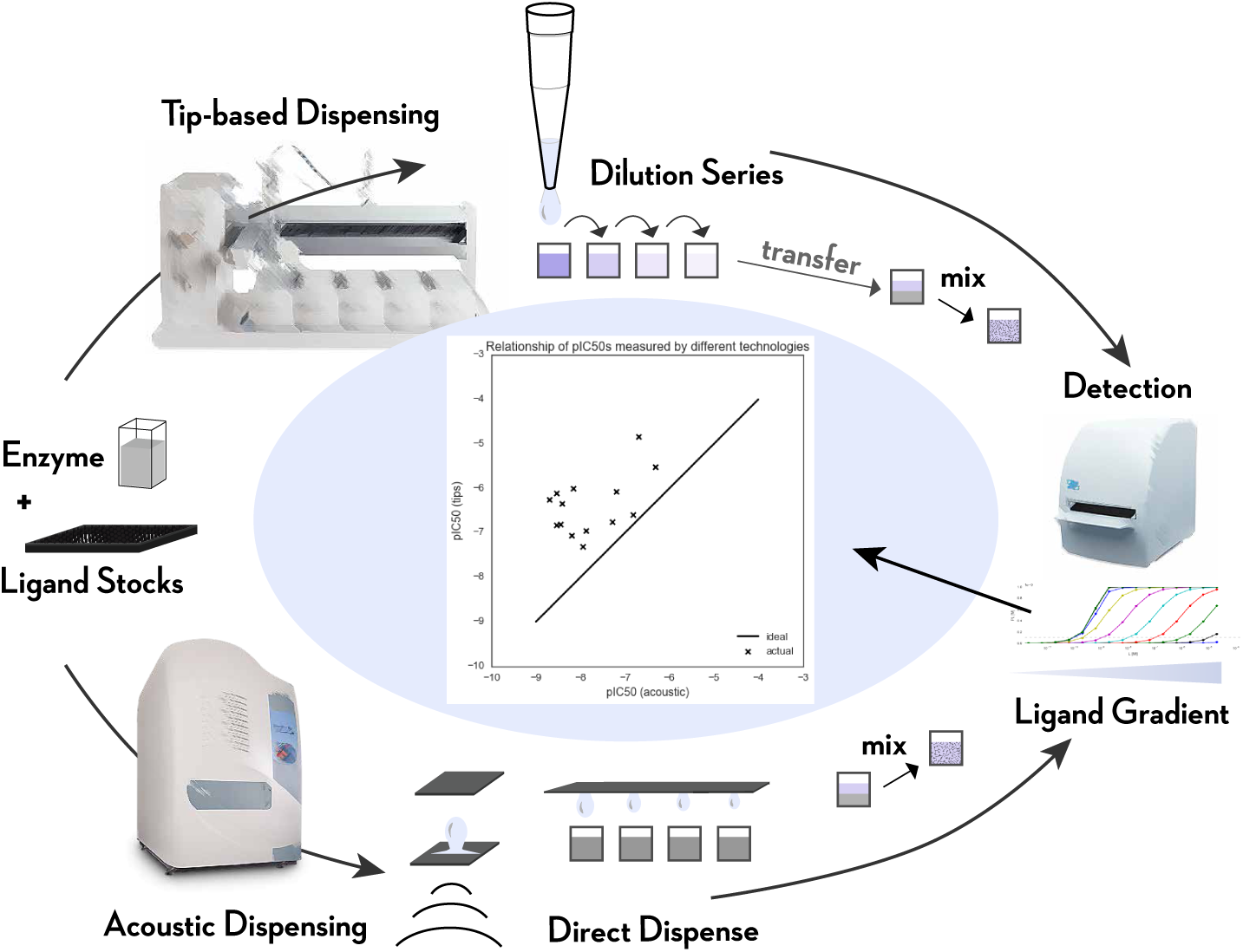
Illustration of the stages of the two assay protocols considered here, utilizing either tip-based or acoustic droplet dispensing. Two different assay protocols—utilizing different dispensing technologies—were used to perform the same assay [19–21]. In the case of tip-based dispensing, a Tecan Genesis liquid handler was used to create a serial dilution of test compounds using fixed washable tips, and a small quantity of each dilution was pipetted into the enzyme assay mixture prior to detection. In the case of acoustic dispensing (sometimes called acoustic droplet ejection), instead of creating a serial dilution, a Labcyte Echo was used to directly dispense nanoliter quantities of compound stock into the enzyme assay mixture prior to detection. The detection phase measured product accumulation after a fixed amount of time (here, detection of accumulated phosphorylated substrate peptide using AlphaScreen), and the resulting data were fit to obtain *p*IC_50_ estimates. Ekins et al. [19] noted that the resulting *p*IC_50_ data between tip-based dispensing and acoustic dispensing were highly discrepant, as shown in the central figure where the two sets of assay data are plotted against each other.

For simplicity, we ask whether the simplest contributions to assay error—imprecision and bias in material transfer operations and imprecision in measurement—might account for some component of the discrepancy between assay techniques. We make use of basic information—the assay protocol as described (with some additional inferences based on fundamental concepts such as compound solubility limits) and manufacturer specifications for imprecision and bias—to construct a model of each dispensing process in order to determine the overall inaccuracy and imprecision of the assay due to dispensing errors, and identify the steps that contribute the largest components to error. To better illustrate these techniques, we also provide an annotated IPython notebook^2^ that includes all of the computations described here in detail. Interested readers are encouraged to download these notebooks and explore them to see how different assay configurations affect assay error, and customize the notebooks for their own scenarios.

## II. EXPERIMENTAL ERROR

Experimental error can be broken into two components: The *imprecision* (quantified by standard deviation or variance), which characterizes the random component of the error that causes different replicates of the same assay to give slightly different results, and the *inaccuracy* (quantified by bias), which is the deviation of the average over many replicates from the true value of the quantity being measured.

There are a wide variety of sources that contribute to experimental error. Variation in the quantity of liquid delivered by a pipette, errors in the reported mass of a dry compound, or noise in the measured detection readout of a well will all contribute to the error of an assay measurement. If the average (mean) of these is the true or desired quantity, then these variations all contribute to imprecision. If not—such as when a calibration error leads to a systematic deviation in the volume delivered by a pipette, the mass measured by a balance, or the average signal measured by a plate reader—the transfers or measurements will also contribute to inaccuracy or bias. We elaborate on these concepts and how to quantify them below.

## MODELING EXPERIMENTAL ERROR

### 1. The hard way: Propagation of error

There are many approaches to the modeling of error and its propagation into derived data. Often, undergraduate laboratory courses provide an introduction to the tracking of measurement imprecision, demonstrating how to propagate imprecision in individual measurements into derived quantities using Taylor series expansions—commonly referred to simply as *propagation of error* [28]. For example, for a function *f* (*x, y*) of two measured quantities *x* and *y* with associated standard errors *σ_x_* and *σ_y_* (which represent our estimate of the standard deviation of repeated measurements of *x* and *y*), the first-order Taylor series error propagation rule is,

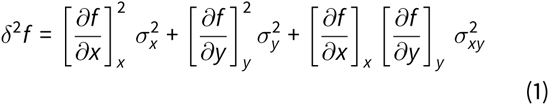

where the correlated error 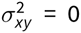 if the measurements of *x* and *y* are independent. The expression for *δ^2^f*, the estimated variance in the computed function *f* over experimental replicates, in principle contains higher-order terms, as well, but first-order Taylor series error propagation presumes these higher-order terms are negligible and all error can be modeled well as a Gaussian (normal) distribution.

For addition or subtraction of two independent quantities, this rule gives a simple, well-known expression for the additivity of errors in quadrature,

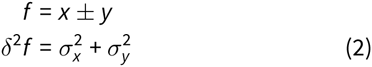

For more complex functions of the data, however, even the simple form of Eq. 1 for just two variables can be a struggle for most scientists to apply, since it involves more complex derivatives that may not easily simplify.

### 2 The easy way: The bootstrap principle

Instead, we adopt a simpler approach based on the *bootstrap principle* [29], *Bootstrapping* allows the sampling distribution to be approximated by simulating from a good estimate (or simulacrum) of the real process, While many computational chemists may be familiar with *resampling boot–strapping* for a large dataset, where resampling values from the dataset with replacement provides a way to simulate a replica of the real process, it is also possible to simulate the process in other ways, such as from a parametric or other model of the process. Here, we model sources of random error using simple statistical distributions, and *simulate* multiple replicates of the experiment, examining the distribution of experimental outcomes in order to quantify error, Unlike propagation of error based on Taylor series approximations (Eq. 1), which can become nightmarishly complex for even simple models, quantifying the error by bootstrap simulation is straightforward even for complex assays. While there are theoretical considerations, practical application of the bootstrap doesn’t even require that the function *f* be differentiable or easily written in closed form—as long as we can compute the function f on a dataset, we can bootstrap it.

For example, for the case of quantities *x* and *y* and associated errors *σ_x_* and *σ_y_*, we would conduct many realizations *n* = 1,…, *N* of an experiment in which we draw *bootstrap replicates x_n_* and *y_n_* from normal (Gaussian) distributions

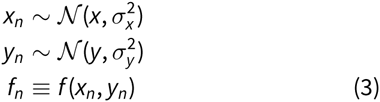

where the notation 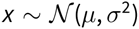 denotes that we draw the variable *x* from a normal (Gaussian) distribution with mean *μ* and variance *σ*^2^,

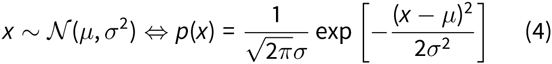

We then analyze the statistics of the {*f_n_*} samples as if we had actually run the experiment many times. For example, we can quantify the statistical uncertainty *δf* using the standard deviation over the bootstrap simulation realizations, std(*f_n_*). Alternatively, presuming we have simulated enough bootstrap replicates, we can estimate 68% or 95% confidence intervals, which may sometimes be very lopsided if the function *f* is highly nonlinear.

Since most instruments we deal with in a laboratory—such as pipettes or liquid handlers or balances—have readily available manufacturer-provided specifications for imprecision and accuracy, we will generally make use of the normal (Gaussian) distribution^3^ in modeling the error Δ*x*,

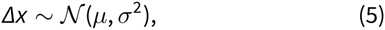

where Δ*x* = *x* − *x*_*_ is the deviation from the true or desired value *x_*_*, the mean σ represents the inaccuracy, and the standard deviation *σ* represents the imprecision.

We will generally quantify the error from our bootstrap simulation replicates in terms of two primary statistics: **Relative bias (RB)**. As a measure of inaccuracy, we will compute the relative expected deviation from the true value *f*,

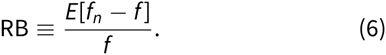

This is often expressed as a percentage (RB%) by multiplying RB by 100. Note that, for cases where *f* = 0, this can be a problematic measure, in which case the absolute bias (just the numerator) is a better choice.

**Coefficient of variation (CV)**. As a measure of imprecision, we will compute the relative standard deviation,

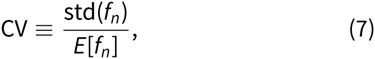

which can again be estimated from the mean over many bootstrap replicates, and is often also represented as a percent (CV%) by multiplying CV by 100.

### Simple liquid handling: Mixing solutions

#### The bootstrap principle

Consider the simplest kind of liquid transfer operation, in which we use some sort of pipetting instrument (hand-held or automated) to combine two solutions. For simplicity, we presume we combine a volume *V*_stock_ of compound stock solution of known true concentration *C*_0_ with a quantity of buffer of volume *V*_buffer_.

Initially, we presume that these operations are free of bias, but have associated imprecisions *σ*_stock_ and *σ*_buffer_. To simulate this process using the bootstrap principle, we simulate a number of realizations *n* = 1,…, *N*, where we again, assume a normal distribution for the sources of error, neglecting bias and accounting only for imprecision,

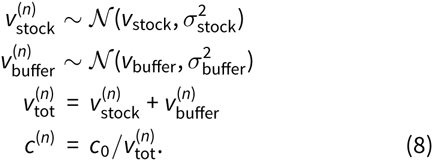

We can then compute statistics over the bootstrap replicates of the resulting solution concentrations, {*c*^(^*^n^*^)^}, *n* = 1,…, *N* to estimate the bias and variance in the concentration in the prepared solution.

#### Relative imprecision

Manufacturer specifications^4^ often provide the imprecision in relative terms as a coefficient of variation (CV), from which we can compute the imprecision σ in terms of transfer volume *v* via *σ* = CV · *v*,

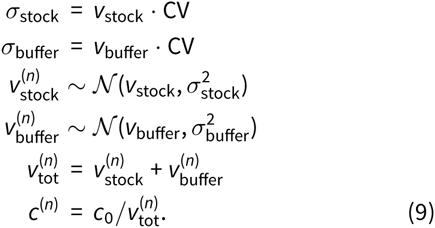

We remind the reader that a CV specified as a % (CV% or %CV) should be divided by 100 to obtain the CV we use here.

#### Relative inaccuracy

Similarly, the expected inaccuracy might also be stated in terms of a relative percentage of the volume being transferred. The inaccuracy behaves differently from the imprecision in that the inaccuracy will *bias* the transferred volumes in a consistent way throughout the whole experiment. To model bias, we draw a single random bias for the instrument from a normal distribution, and assume all subsequent operations with this instrument are biased in the same relative way. We presume the relative bias (RB)–expressed as a fraction, rather than a percent—is given as RB, and draw a specific instrumental bias *b*^(^*^n^*^)^ for each bootstrap replicate of the experiment, simulating the effect of many replications of the experiment where the instrument is randomly recalibrated,

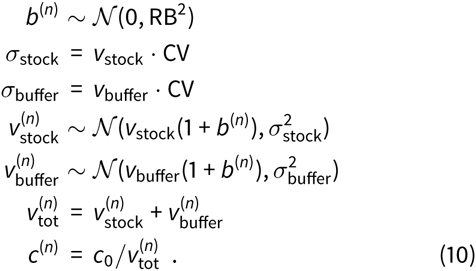

Obviously, if the instrument is never recalibrated, the bias will be the *same* over many realizations of the experiment, but we presume that a calibration process is repeated frequently enough that its effect can be incorporated as a random effect over many replicates of the assay over a long timespan.

#### Uncertainty in initial concentration

We could further extend this model to include uncertainty *σ_c_* in the stock concentration *c*_0_ (where the concentration may be stated *c*_0_ ± *σ_c_*), and begin to see how powerful and modular the bootstrap scheme is. In this new model, each simulation realization n includes one additional initial step,

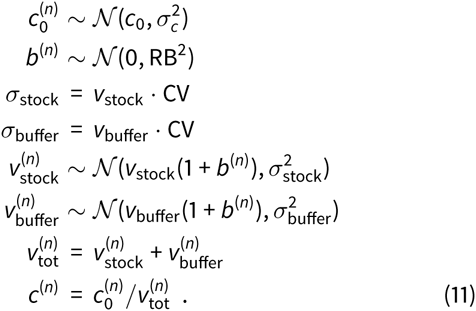

All we had to do was add one additional step to our bootstrap simulation scheme (Eq. 10) in which the stock concentration 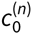 is independently drawn from a normal distribution with each bootstrap realization *n*. The model can be expanded indefinitely with additional independent measurements or random variables in the same simple way.

Below, we exploit the modularity of bootstrap simulations to design a simple scheme to model a real assay—the measurement of *p*IC_50_s for compounds targeting the EphB4 receptor [19–21]—without being overwhelmed by complexity. This assay is particularly interesting because data exists for the same assay performed using two *different* dispensing protocols that led to highly discrepant assay *p*IC_50_ data, allowing us to examine how different sources of error arising from different dispensing technologies can impact an otherwise identical assay. We consider only errors that arise from the transfer and mixing of volumes of solutions with different concentrations of compound, using the same basic strategy seen here to model the mixing of two solutions applied to the complex liquid handling operations in the assay. To more clearly illustrate the impact of imprecision and inaccuracy of dispensing technologies, we neglect considerations of the completeness of mixing, which can itself be a large source of error in certain assays^5^.

### Modeling an enzymatic reaction and detection of product accumulation

The EphB4 assay we consider here [19–21], illustrated schematically in Figure 1, measures the rate of substrate phosphorylation in the presence of different inhibitor concentrations. After mixing the enzyme with substrate and inhibitor, the reaction is allowed to progress for one hour before being quenched by the addition of a quench buffer containing EDTA. The assay readout (in this case, AlphaScreen) measures the accumulation of phosphorylated substrate peptide. Fitting a binding model to the assay readout over the range of assayed inhibitor concentrations yields an observed *p*IC_50_.

A simple model of inhibitor binding and product accumulation for this competition assay can be created using standard models for competitive inhibition of a substrate S with an inhibitor /. Here, we assume that in excess of substrate, the total accumulation of product in a fixed assay time will be proportional to the relative enzyme turnover velocity times time, *V*_0_*t*, and use an equation derived assuming Michaelis-Menten kinetics,

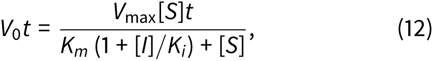

where the Michaelis constant *K_m_* and substrate concentration [S] forthe EphB4 system are pulled directly from the assay methodology description [20, 21]. To simplify our modeling, we divide by the constants *V*_max_*t*, and work with the simpler ratio,

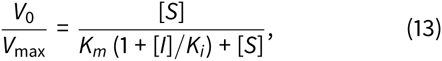

In interrogating our model, we will vary the true inhibitor affinity *K_i_* to determine how the assay imprecision and inaccuracy depend on true inhibitor affinity.

In reality, detection of accumulated product will also introduce uncertainty. First, there is a minimal detectable signal below which the signal cannot be accurately quantified—below this threshold, a random background signal or “noise floor” is observed. Second, any measurement will be contaminated with noise, though changes to the measurement protocol—such as collecting more illumination data at the expense of longer measurement times—can affect this noise. While simple calibration experiments can often furnish all of the necessary parameters for a useful detection error model—such as measuring the background signal and signal relative to a standard for which manufacturer specifications are available—we omit these effects to focus on the potential for the discrepancy between liquid handling technologies to explain the difference in assay results.

### Advanced liquid handling: Making a dilution series

Because the affinities and activities of compounds can vary across a dynamic range that spans several orders of magnitude, it is common for assays to use a dilution series to measure the activity and potency of ligands. To create a dilution series, an initial compound stock is diluted into buffer in the first well of the series, and the contents mixed; for each subsequent well, a volume from the previous well is transferred into a well containing only buffer, and mixed (Figure 2). Commonly, each subsequent dilution step uses fixed ratios, such as 1:2 or 1:10 of solute solution to total volume^6^.

**FIG. 2.**
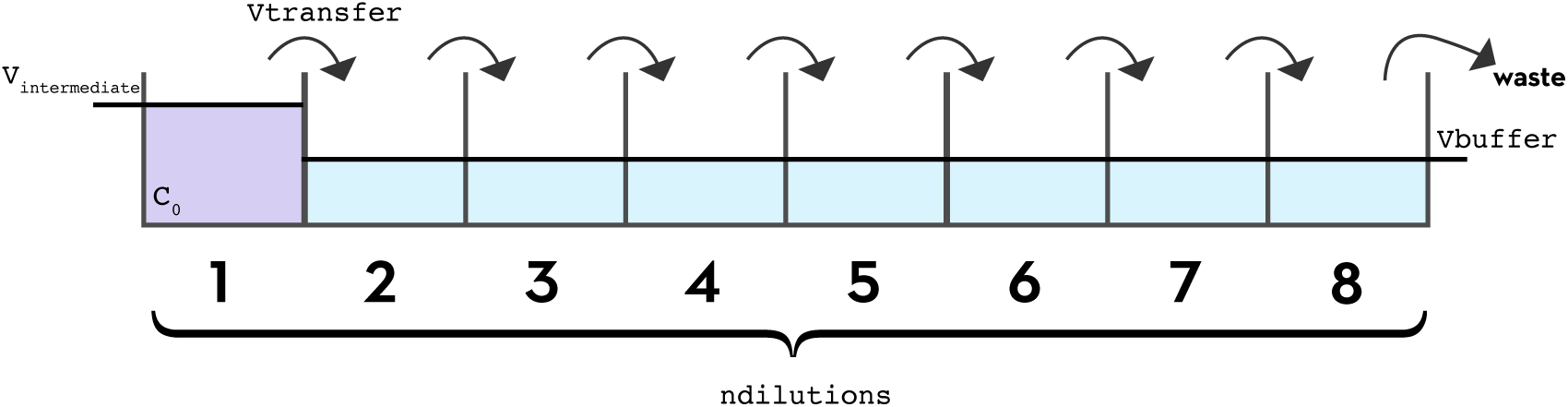
Preparation of a serial dilution series with a fixed-tip liquid handler. To create a dilution series with a fixed-tip liquid handler, a protocol similar to the preparation of a dilution series by hand pipetting is followed. Starting with an initial concentration *c*_0_ and initial volume *V*_initial_ in the first well, a volume *v*_transfer_ is transferred from each well into the next well, in which a volume of buffer, *V*_buffer_, has already been pipetted. In the case of a 1:2 dilution series, *v*_transfer_ and *v*_buffer_ are equal, so the intended concentration in the second well will be *c*_0_/2. This transfer is repeated for all subsequent wells to create a total of *n*_dilutions_ dilutions. For convenience, we assume that a volume *v*_transfer_ is removed from the last well so that all wells have the same final volume of *v*_transfer_ = *v*_buffer_, and that the error in the initial concentration (*c*_0_) is negligible.

It is easy to see how the creation of a dilution series by pipetting can amplify errors: because each dilution step involves multiple pipetting operations, and the previous dilution in the series is used to prepare the next dilution, errors will generally grow with each step. As a result, the liquid handling instrumentation can have a substantial impact on the results obtained. Here, we compare an aqueous dilution series made with a liquid handler that makes use of fixed, washable tips (a Tecan Genesis) with an assay prepared directly via direct-dispensing using an acoustic dispensing instrument (a Labcyte Echo).

#### Tip-based liquid handling

To create a serial dilution series (Figure 2), we first transfer an aliquot of compound in DMSO stock solution to the first well, mixing it with buffer, to prepare the desired initial concentration *c*_0_ at volume *v*_intermediate_ for the dilution series. Next, we sequentially dilute a volume *V*_transfer_ of this solution with a volume *V*_buffer_ of buffer, repeating this process to create a total of *n*_dilutions_ solutions. To model the impact of imprecision and inaccuracy on the serial dilution process, we again use manufacturer-provided specifications for the Tecan Genesis: the relative imprecision is stated to be 3% and the inaccuracy as 3–5% for the volumes in question [33]. The resulting concentration *c_m_* of each dilution *m* is determined by both the previous concentration *c_m_*_−1_ and by the pipetted volumes *V*_transfer_ and *V*_buffer_, each of which is randomly drawn from a normal distribution. Putting this together in the same manner as for the simple mixing of solutions, we have

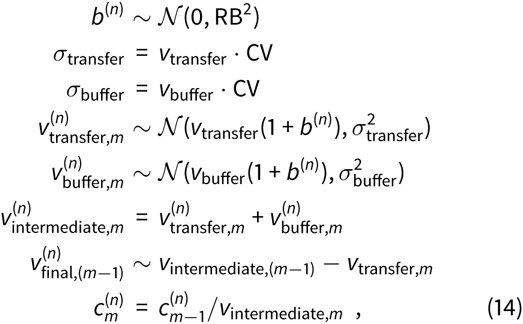

where the last five steps are computed for dilution *m* = 1,…, (*n*_dilutions_ − 1). In the companion notebook, we make comparisons easier by also removing a final volume *v*_transfer_ from the last well so all wells have the same intended final volume.

In the EphB4 protocol [20, 21], the initial dilution step from 10 mM DMSO stocks is not specified, so we choose initial concentration *c*_0_ = 600 *μ*M in order to match the maximum assay concentration used in the direct dispense version of the assay (described in the next section). We presume an initial working volume of *v*_intermediate_ = 100 *μ*L, and model this dispensing process using Eq. 10. We presume this solution is then serially 1:2 diluted with 5% DMSO for a total of *n*_dilutions_ = 8 dilutions^7^ with *V*_transfer_ = *V*_buffer_ = 50 *μ*L, which after dilution into the assay plate will produce a range of assay concentrations from 800 nM to 100 *μ*M^8^. We estimate the appropriate coefficient of variation (CV) and relative bias (RB) for the Tecan Genesis liquid handling instrument used in this assay using a linear interpolation over the range of volumes in a manufacturer-provided table [33]. Sampling over many bootstrap replicates, we are then able to estimate the CV and RB in the resulting solution concentrations for the dilution series. Figure 5 (left panel) shows the estimated CV and RB for the resulting concentrations in the dilution series. While the CV for the volume is relatively constant since we are always combining only two transferred aliquots of liquid, the CV for both the concentration and the total quantity of compound per well grow monotonically with each subsequent dilution. On the other hand, because the bias is assumed to be zero on average, the average dilution series bias over many bootstrap replicates with randomly calibrated instruments will be free of bias. This situation may be different, of course, if the *same* miscalibrated instrument is used repeatedly without frequent recalibration.

Once the dilution series has been prepared, the assay is performed in a 384-well plate, with each well containing 2 *μ*L of the diluted compound in buffer combined with 10 *μ*L of assay mix (which contains EphB4, substrate peptide, and cofactors) for a total assay volume of 12 *μ*L. This liquid transfer step is easily modeled using the steps for modeling the mixing of two solutions in Eq. 10.

#### Direct dispensing technologies

Using a direct dispensing technology such as acoustic droplet ejection (ADE), we can eliminate the need to prepare an intermediate dilution series, instead addingsmall quantities of the compound DMSO stock solution directly into the assay plate. For the LabCyte Echo used in the EphB4 assay, the smallest volume dispensed is 2.5 nL droplets; other instruments such as the HP D300/D300e can dispense quantities as small as 11 pL using inkjet technology. To construct a model for a direct dispensing process, we transfer a volume *v*_dispense_ of ligand stock in DMSO at concentration *c*_0_ into each well already containing assay mix at volume *v*_mix_ (presumed to be pipetted by the Tecan Genesis), and backfill a volume *v*_backfill_ with DMSO to ensure each well has the same intended volume and DMSO concentration (Figure 3). We again incorporate the effects of imprecision and bias using manufacturer-provided values; for the Labcyte Echo, the relative imprecision (CV) is stated as 8% and the relative inaccuracy (RB) as 10% for the volumes in question [34],

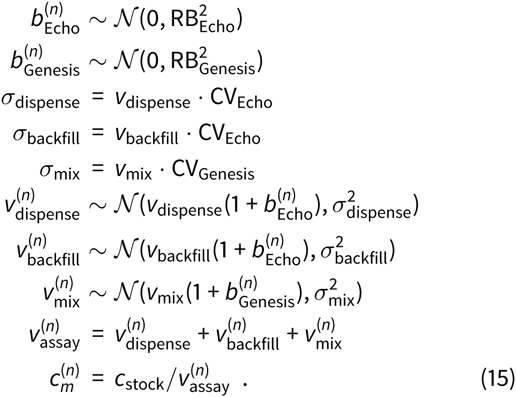

**FIG. 3.**
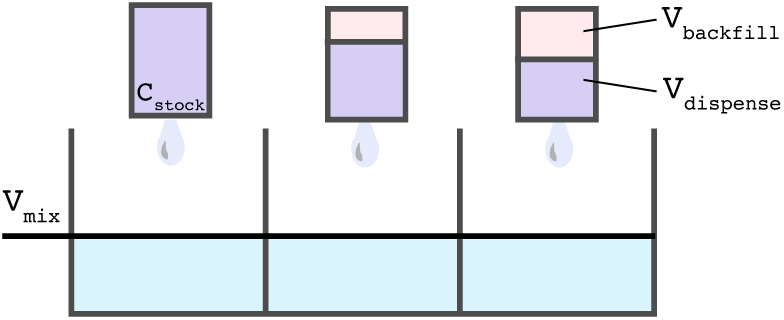
Preparation of a dilution series with direct-dispense technology. With a direct-dispense liquid handler—such as the LabCyte Echo, which uses acoustic droplet ejection—instead of first preparing a set of compound solutions at different concentrations via serial dilution, the intended quantity of compound can be dispensed into the assay plates directly without the need for creating an intermediate serial dilution. We model this process by considering the process of dispensing into each well independently. A volume *V*_dispense_ of compound stock in DMSO at concentration *c*_0_ is dispensed directly into an assay plate containing a volume *V*_mix_ of assay mix. To maintain a constant DMSO concentration throughout the assay—in this case of the EphB4 assay, 120 nL—a volume *V*_backfill_ of pure DMSO is also dispensed via acoustic ejection.

Since the maximum specified backfilled volume was 120 nL, we presume that *v*_dispense_ consisted of 8 dilutions ranging from 2.5 nL (the minimum volume the Echo can dispense) to 120 nL in a roughly logarithmic series. Note that this produces a much narrower dynamic range than the dilution series experiment, with the minimum assay intended concentration being 2.5 *μ*M assuming a 10 mM DMSO stock solution concentration *c*_stock_. We can then produce an estimate for the errors in volumes and concentrations (Figure 5, middle panels) by generating many synthetic replicates of the experiment. Because direct dispensing technologies can dispense directly into the assay plate, rather than creating an intermediate dilution series that is then transferred into the assay wells, direct dispensing experiments can utilize fewer steps (and hence fewer potential inaccuracy- and imprecision-amplifying steps) than the tip-based assays that are dependent on the creation of an intermediate dilution series.

Simply including the computed contributions from inaccuracy and imprecision in our model of the Ekins et al. dataset [19], it is easy to see that the imprecision is not nearly large enough to explain the discrepancies between measurements made with the two dispensing technologies (Figure 7). Multichannel liquid-handlers such as the Tecan Genesis that utilize liquid-displacement pipetting with fixed tips actually have a nonzero bias in liquid transfer operations due to a dilution effect. This effect was previously characterized in work from Bristol Myers Squibb (BMS) [35, 36], where it was found that residual system liquid—the liquid used to create the pressure differences required for pipetting—can cling to the interior of the tips after washing and mix with sample when it is being aspirated (Figure 4). While the instrument can be calibrated to dispense volume without bias, the concentration of the dispensed solution can be measurably diluted.

**FIG. 4.**
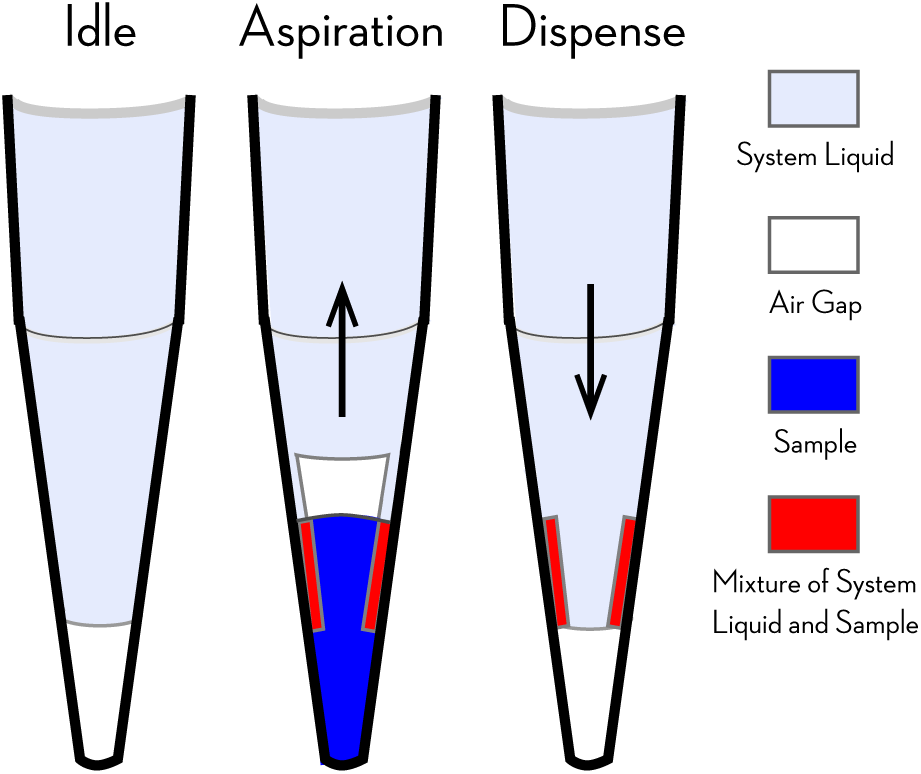
Fixed tips dilute aspirated samples with system liquid. Automated liquid handlers with fixed tips utilizing liquiddisplacement pipetting technology (such as the Tecan Genesis used in the EphB4 assay described here) use a washing cycle in which system liquid (generally water or buffer) purges samples from the tips in between liquid transfer steps. Aspirated sample (blue) can be diluted by the system liquid (light purple) when some residual system liquid remains wetting the inside walls of the tip after purging. This residual system liquid is mixed with the sample as it is aspirated, creating a mixture of system liquid and sample (red) that dilutes the sample that is dispensed. While the use of an air gap (white) reduces the magnitude of this dilution effect, dilution is a known issue in fixed tip liquid-based automated liquid handlingtechnologies, requiring more complex liquid-handling strategies to eliminate it [36]. Diagram adapted from Ref. [36].

To quantify this effect, the BMS team used both an Artel dye-based Multichannel Verification System (MVS) and gravimetric methods, concluding that this dilution effect contributes a −6.30% inaccuracy for a target volume of 20 *μ*L [35]. We can expand our bootstrap model of dilution with fixed tips (Eq. 15) to include this effect with a simple modification to the concentration of dilution solution *m*,

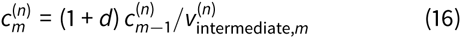

where the factor *d* = −0.0630 accounts for the −6.30% dilution effect. The resulting CV and RB in volumes, concentrations, and quantities (Figure 5, middle) indicate a significant accumulation of bias. This is especially striking when considered alongside the corresponding values for disposable tips (Figure 5, left)—which lack the dilution effect—and the acoustic-dispensing model (Figure 5, right), both of which are essentially free of bias when the average over many random instrument recalibrations is considered.

**FIG. 5.**
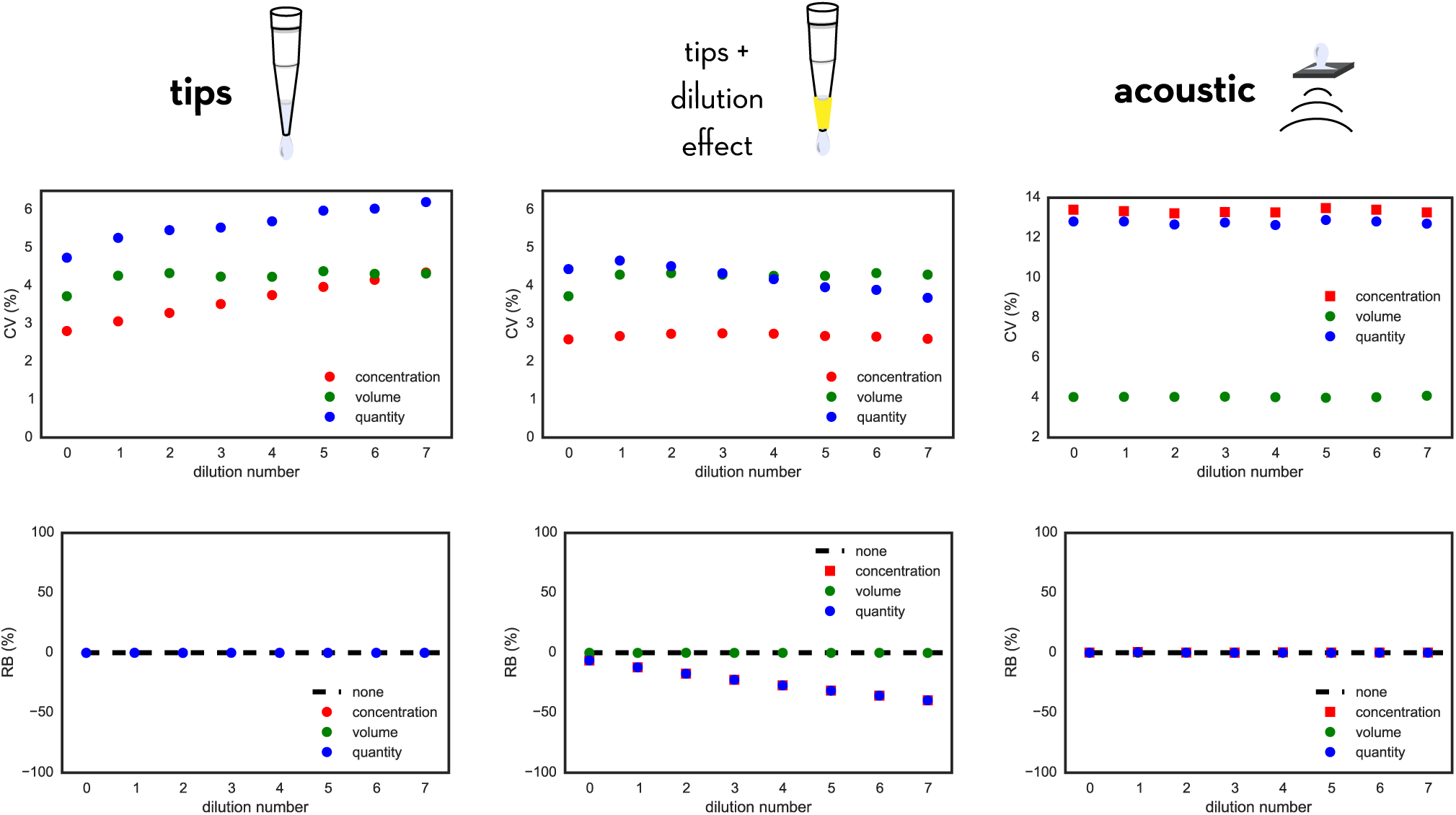
Modeled accumulation of random and systematic error in creating dilution series with fixed tips and acoustic dispensing. The model predicts how errors in compound concentration, well volume, and compound quantity accumulate for a dilution series prepared using fixed tips neglecting dilution effects *(left)* or including dilution effects *(middle)* compared with an acoustic direct-dispensing process *(right)*. Imprecision and inaccuracy parameters appropriate for a Tecan Genesis (fixed tips dispensing) or Labcyte Echo (acoustic dispensing) were used, and assume that the initial compound stocks had negligible concentration error; see text for more details. The top panels show the average relative random error via the coefficient of variation (CV) of concentration, volume, or quantity, while the bottom panels depict the relative bias (RB); both quantities are expressed as a percentage. For tip-based dispensing, relative random concentration error (CV) accumulates with dilution number, while for acoustic dispensing, this is constant over all dilutions. When the dilution effect is included for fixed tips, there is significant bias accumulation over the dilution series. Note that the CV and RB shown for acoustic dispensing are for the final assay solutions, since no intermediate dilution series is created.

This dilution effect also must be incorporated into the transfer of the diluted compound solutions (2 *μ*L) into the enzyme assay mix (10 *μ*L) to prepare the final 12 *μ*L assay volume, further adding to the overall bias of the assay results from the fixed-tips instrument.

### Fitting the assay readout to obtain *p*IC_50_ data

While the IC_50_ reported in the EphB4 assay [19–21] in principle represents the stated concentration of compound required to inhibit enzyme activity by half, this value is estimated in practice by numerically fitting a model of inhibition to the measured assay readout across the whole range of concentrations measured using a method such as least-squared (the topic of another article in this series [37]).

To mimic the approach used in fitting the assay data, we use a nonlinear least-squares approach (based on the simple `curve_fit` function from `scipy.optimize`) to fit *V*_0_/*V*_max_ computed from the competitive inhibition model (Eq. 13, shown in Fig. 6, top panels) using the true assay well concentrations to obtain a *K_i_* and then compute the IC_50_ from this fit value. We can then use a simple relation between IC_50_ and *K_i_* to compute the reported assay readout,

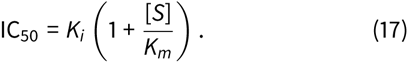

**FIG. 6.**
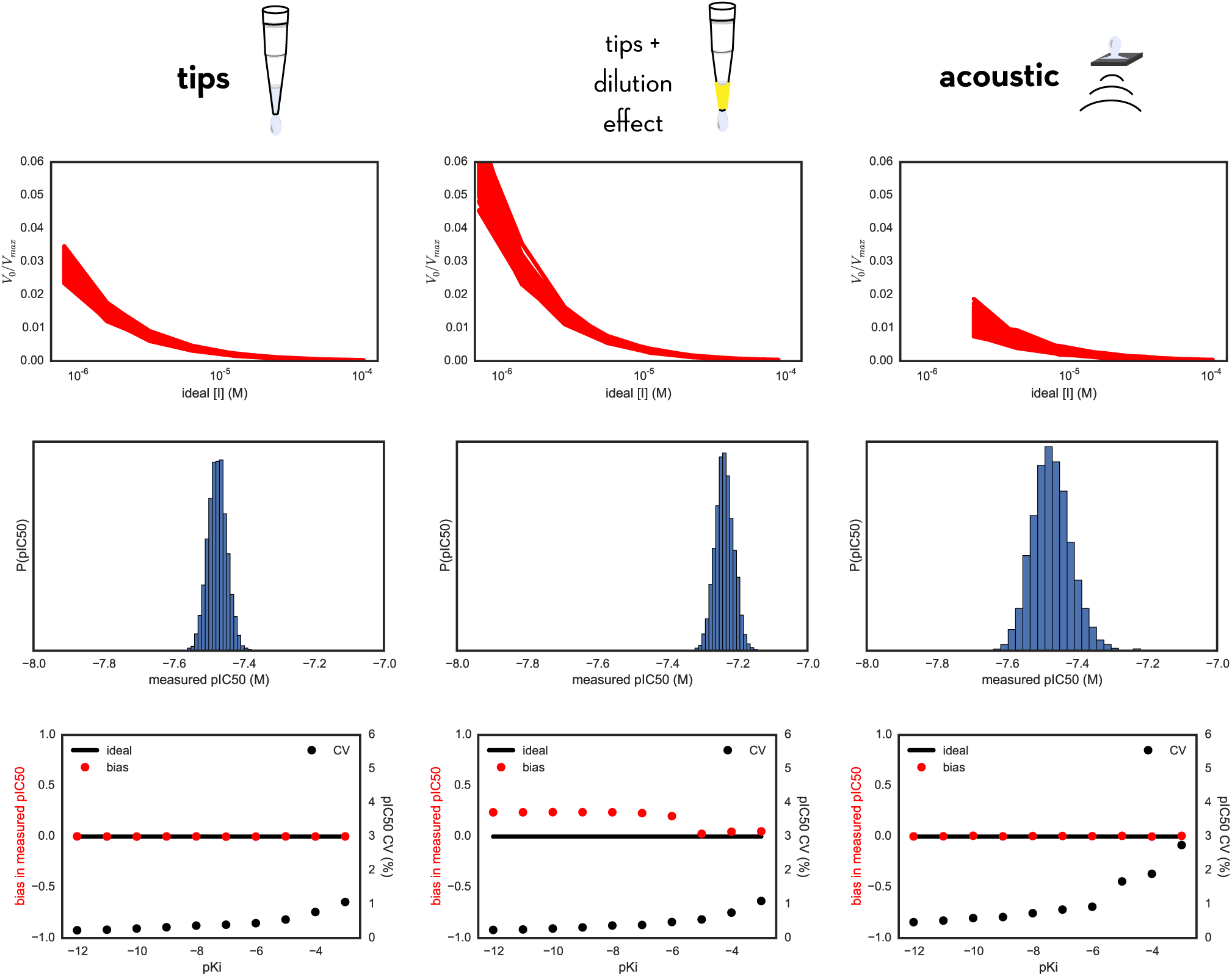
Comparing modeled errors in measured *p*IC_50_ values using tip-based or acoustic direct dispensing. *Top row:* Bootstrap simulation of the entire assay yields a distribution of *V*_0_/*V*_max_ (proportional to measured product accumulation) vs ideal inhibitor concentration [*I*] curves for many synthetic bootstrap replicates of the assay. Here, the inhibitor is modeled to have a true *K_i_*, of 1 nM (*pK_i_* = −9). *Middle row:* For the same inhibitor, we obtain a distribution of measured *p*IC_50_ values from fitting the Using our model we can look at the variance in activity measurements as a function of inhibitor concentration [*I*] (top), which then directly translates into a distribution of measured *p*IC_50_ values. *Bottom row:* Scaning across a range of true compound affinities, we can repeat the bootstrap sampling procedure and analyze the distribution of measured *p*IC_50_ values to obtain estimates of the relative bias (red) and CV (black) for the resulting measured *p*IC_50_s. For all methods, the CV increases for weaker affinities; for tip-based dispensing using fixed tips and incorporating the dilution effect, a significant bias is notable.

The reported results are not IC_50_ values but *p*IC_50_ values,

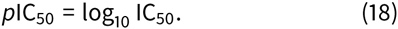

Note that no complicated manipulation of these equations is required. As can be seen in the companion IPython notebook, we can simply use the `curve_fit` function to obtain a *K_i_* for each bootstrap replicate, and then store the *p*IC_50_ obtained from the use of Eqs. 17 and 18 above (Fig. 6, middle panels). Repeating this process for a variety of true compound affinities allows the imprecision (CV) and bias (RB) to be quantified as a function of true compound affinity (Fig. 6, bottom panels).

## III. DISCUSSION

### Use of fixed washable tips can cause significant accumulation of bias due to dilution effects

The most striking feature of Fig. 5 is the significant accumulation of bias in the preparation of a dilution series using fixed washable tips (Fig. 5, bottom middle panel). Even for an 8-point dilution series, the relative bias (RB) is almost −50% in the final well of the dilution series. As a result, the measured *p*IC_50_ values also contain significant bias toward weaker affinities (Fig. 6, bottom middle panel) by about 0.25 log_10_ units for a large range of compound affinities. At weaker compound affinities, this effect is diminished by virtue of the fact that the first few wells of the dilution series have a much smaller RB (Fig. 5, bottom middle panel).

This cumulative dilution effect becomes more drastic if the dilution series is extended beyond 8 points. If instead a dilution series is created across 16 or 32 wells and assayed, the RB in the final well of the dilution series can reach nearly −100% (see accompanying IPython notebook for 32-well dilution series). As a result, the bias in the measured *p*IC_50_ as a function of true *pK_i_*, also grows significantly for these larger dilution series (Fig. 8).

### Imprecision is greater for direct dispensing with the Echo

As evident from the top panels of Fig. 5, the CV for concentrations in the assay volume for direct acoustic dispensing (right) is significantly higher than the CV of the dilution series prepared with tips (left and middle). This effect manifests itself in the CV of measured *p*IC_50_ values as a higher imprecision (Fig. 6, bottom panels), where the CV for acoustic dispensing is nearly twice that of tip-based dispensing. Despite the increased CV, there are still numerous advantages to the use of direct dispensing technology: Here, we have ignored a number of difficulties in the creation of a dilution series beyond this dilution effect, including the difficulty of attaining good mixing [30–32], the time required to prepare the serial dilution series (during which evaporation may be problematic), and a host of other issues.

### Imprecision is insufficient to explain the discrepancy between assay technologies

Fig. 7 depicts the reported assay results [19–21] augmented with error bars and corrected for bias using models appropriate for disposable tips (blue circles) or fixed washable tips (green circles) that include the dilution effect described in Fig. 4. Perfect concordance of measured *p*IC_50_s between assay technologies would mean all points fall on the black diagonal line. We can see that simply adding the imprecision in a model with fixed tips (Fig. 4, blue circles, horizontal and vertical bars denote 95% confidence intervals) is insufficient to explain the departure of the dataset from this diagonal concordance line.

**FIG. 7.**
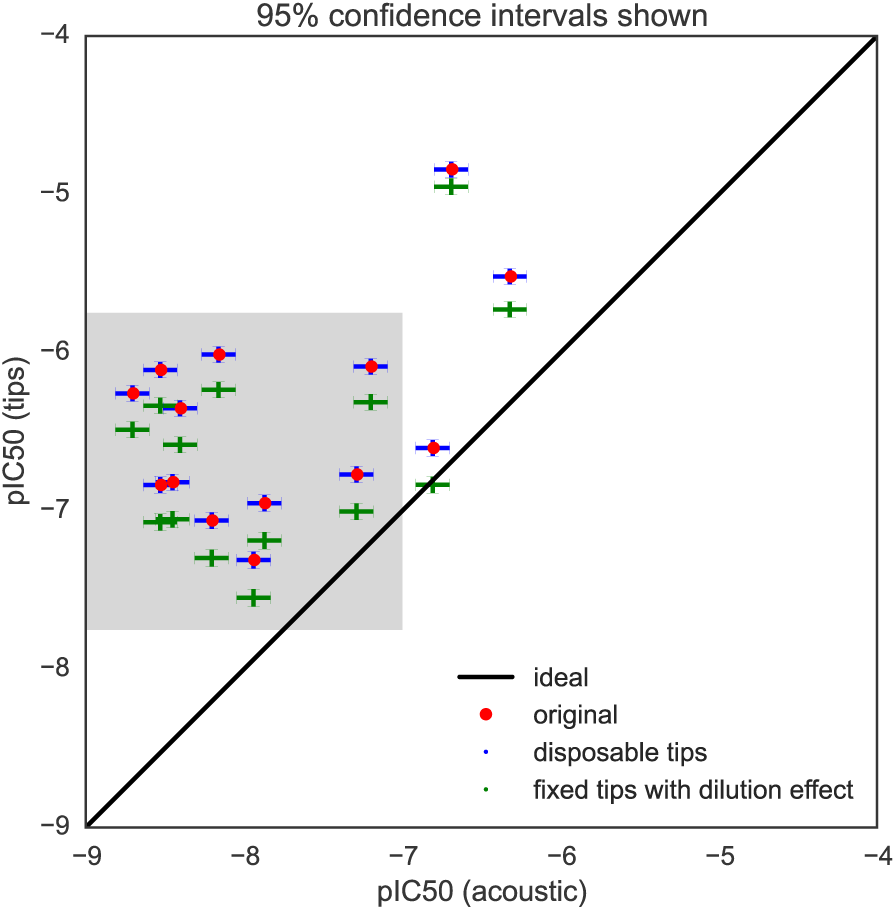
Adding bias shifts *p*IC_50_ values closer to equivalence. The original experimental pIC50 values obtained using from fixed tips (red) are plotted against *p*IC_50_ values from acoustic dispensing, with errors bars representing the uncertainty (shown as 95% confidence intervals) estimated by bootstrapping from our models. Since the bias is relatively sensitive to *p*IC50 value, here it is determined by including both the experimental value and the estimated bias. Incorporating the dilution effect from tip-based dispensing (green) shifts the experimental *p*IC_50_ values closer to concordance between tip-based and acoustic-based measurements. While this does not entirely explain all discrepancies between the two sets of data, it shifts the root mean square error between the tip-based and acoustic-based dispensing methods from 1.56 to 1.37 *p*IC50 units. The model also demonstrates that (1) the bias induced by the fixed tips explains much of the *p*IC_50_ shift between the two datasets, and (2) there is still a large degree of variation among the measurements not accounted for by simple imprecision in liquid transfers. This demonstrates the power of building simple error models to improve our understanding of experimental data sets. Grey box indicates portion of graph shown in Fig. 8.

When the tip dilution effect for washable tips is incorporated (Fig. 4, green circles), there is a substantial shift toward higher concordance. If, instead of an 8-point dilution series, a 16- or 32-point dilution series was used, this shift toward concordance is even larger (Fig. 8). While this effect may explain a substantial component of the divergence between assay technologies, there is no doubt a significant discrepancy remains.

**FIG. 8.**
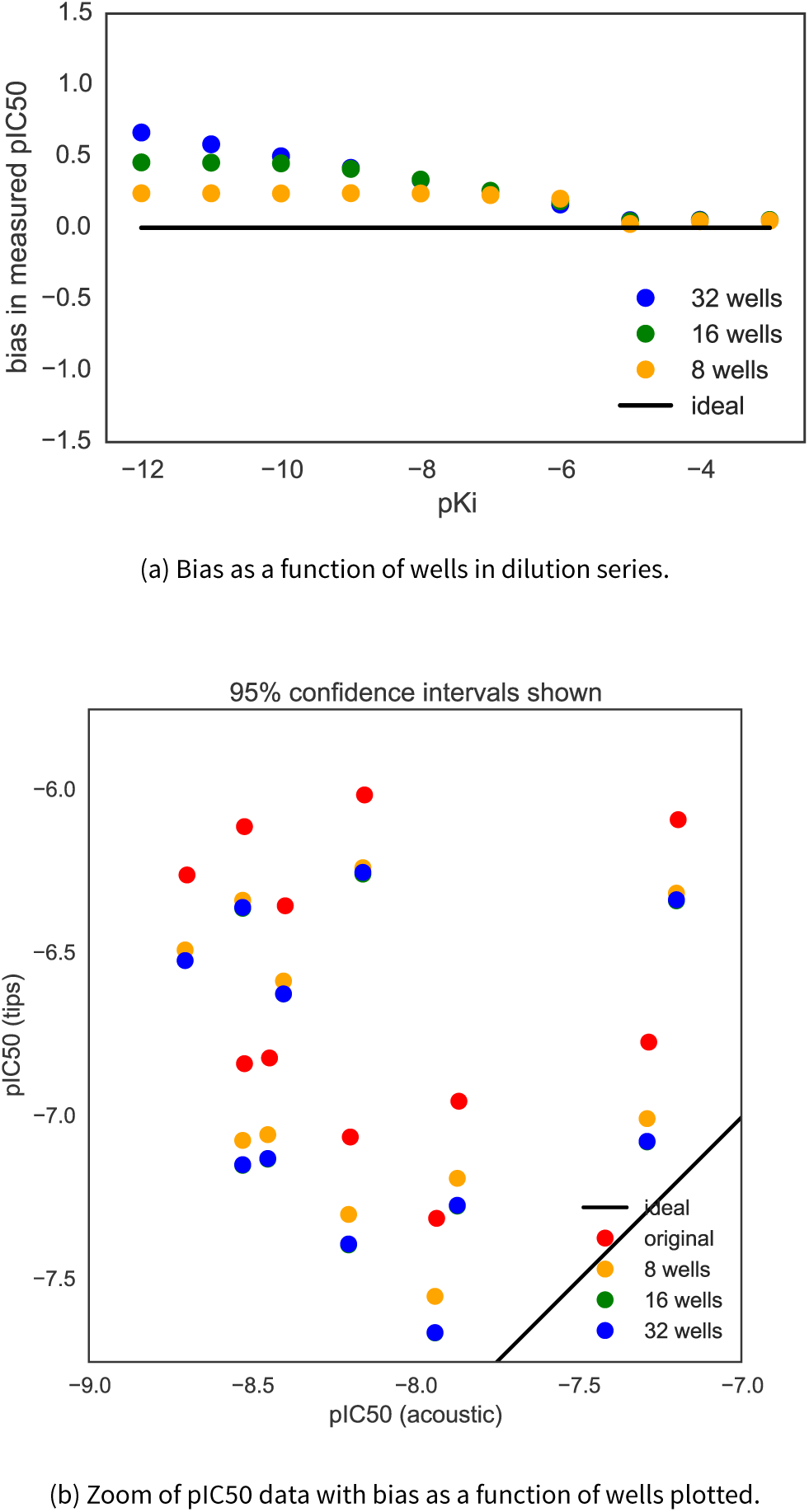
Bias in measured *p*IC_50_ depends on number of wells in dilution series when using fixed washable tips. (a) If the dilution series is extended beyond 8 wells (yellow) to instead span 16 (green) or 32 (blue) wells, the bias effect in the measured pIC_50_ increases as the cumulative effect of the dilution effect illustrated in Fig. 4 shifts the apparent affinity of the compound. Because the dilution bias is greater for lower compound concentrations, this effect is more drastic for compounds with high affinity. (b) Applying these biases to the *p*IC50 from the sample dataset shows the bias increases with both the 16 (green) and 32 (blue) well dilution series, shifting the points even further toward the line of ideal equivalence of the two types of liquid handling. Note these points overlap exactly.

### Other contributions to the discrepancy are likely relevant

Serial dilutions are commonly used in the process of determining biologically and clinically relevant values such as inhibition concentrations (IC_50_) and dissociation constants (*K_d_*). While high-throughput automation methods can improve the reproducibility of these measurements over manual pipetting, even robotic liquid handlers are victim to the accumulation of both random and systematic error. Since the AstraZeneca dataset [20, 21] and the related analysis by Ekins et al. [19], several studies have posited that acoustic dispensing results in fewer false positives and negatives than tip-based dispensing and that this phenomenon is not isolated to EphB4 receptor inhibitors [38–41].

### The power of bootstrapping

We have demonstrated how a simple model based on the bootstrap principle, in which nothing more than the manufacturer-provided imprecision and inaccuracy values and a description of the experimental protocol were used to *simulate* virtual replicates of the experiment for a variety of simulated compound affinities allowed us to estimate the imprecision and inaccuracy of measured *p*IC_50_s. It also identified the difficulty in creating an accurate dilution series using washable fixed tips, with the corresponding dilution effect being a significant contribution to discrepancies in measurements between fixed pipette tips and direct dispensing technologies. In addition to providing some estimate for the random error in measured affinities, the computed bias can even be used to *correct* for the bias introduced by this process after the fact, though it is always safer to take steps to minimize this bias before the assay is performed.

The EphB4 assay considered here is just one example of a large class of assays involving dilution or direct dispensing of query compounds followed by detection of some readout. The corresponding bootstrap model can be used as a template for other types of experiments relevant to computational modelers.

This approach can be a useful general tool for both experimental and computational chemists to understand common sources of error within assays that use dilution series and how to model and correct for them. Instead of simply relying on intuition or historically successful protocol designs, experimentalists could use bootstrap simulation models during assay planning stages to verify that the proposed assay protocol is capable of appropriately discriminating among the properties of the molecules in question given the expected range of IC_50_ or K*_i_* to be probed, once known errors are accounted for. Since the model is quantitative, adjusting the parameters in the assay protocol could allow the experimentalist to optimize the protocol to make sure the data is appropriate to the question at hand. For example, in our own laboratory, it has informed the decision to use only direct dispensing technologies—in particular the HP D300 [42]—for fluorescent ligand-binding assays that require preparation of a range of compound concentrations.

This modeling approach can also be extremely useful in determining appropriate tests and controls to use to be sure errors and biases are properly taken into account in general. If one is not certain about the primary sources of error in an experiment, one is arguably not certain about the results of the experiment in general. Understanding these errors, and being certain they are accounted for via clear benchmarks in experimental assays could help ensure the reproducibility of assays in the future, which is currently a topic of great interest. Especially with such a wide ranging set of assays that use dilution series, most notably toward the development of proteins and small molecules to study and treat disease, this is a very important category of experiments to understand how to make more clearly reproducible and interpretable.

While here we have illustrated the importance of modeling to the specific case of liquid handling with fixed tips in the context of measuring IC_50_ values for EphB4 inhibitors, there are still large discrepancies that have not been explained, and perhaps variations on this model could explain everything, but perhaps the full explanation comes from parts of the assay yet to be incorporated into this model. As experiments become more automated and analysis becomes more quantitative, understanding these errors will be increasingly important both for the consumers (computational modelers) and producers (experimentalists) of these data.

## IV. ACKNOWLEDGMENTS

The authors are grateful to Anthony Nichols (OpenEye) and Martin Stahl (Roche) for organizing the excellent 2013 Computer-Aided Drug Discovery Gordon Research Conference on the topic of “The Statistics of Drug Discovery”, as well as Terry Stouch for both his infinite patience and inspiring many of the ideas in this work. The authors are especially grateful to Cosma Shalizi for presenting a clear and lucid overview of the bootstrap principle to this audience, and we hope this contribution can further aid readers in the community in employing these principles in their work. The authors further acknowledge Adrienne Chow and Anthony Lozada of Tecan US for a great deal of assistance in understanding the nature of operation and origin of errors in automated liquid handling equipment. The authors thank Paul Czodrowski (Merck Darmstadt) for introducing us to IPython notebooks as a means of interactive knowledge transfer. JDC and SMH acknowledge support from the Sloan Kettering Institute, a Louis V. Gerstner Young Investigator Award, and NIH grant P30 CA008748. SE acknowledges Joe Olechno and Antony Williams for extensive discussions on the topic, as well as the many scientists that responded to the various blog posts mentioned herein.

## Footnotes

1 Care must be taken to distinguish between fully independent replicates and partial replicates that only repeat part of the experiment (for example, repeated measurements performed using the same stock solutions), since partial measurements can often underestimate true error by orders of magnitude [16].

2 The companion IPython notebook is available online at: http://github.com/choderalab/dispensing-errors-manuscript

3 Volumes, masses, and concentrations must all be positive, so it is more appropriate in principle to use a *lognormal* distribution to model these processes to prevent negative values. In practice, however, if the relative imprecision is relatively small and negative numbers do not cause large problems for the functions, a normal distribution is sufficient.

4 While manufacturer-provided specifications for imprecision and inaccuracy are often presented as the maximum-allowable values, we find these are a reasonable starting point for this kind of modeling.

5 A surprising amount of effort is required to ensure thorough mixing of two solutions, especially in the preparation of dilution series [30–32]. We have chosen not to explicitly include this effect in our model, but it could similarly be added within this framework given some elementary data quantifying the bias induced by incomplete mixing.

6 Note that a 1:2 dilution refers to combining one part solute solution with one part diluent.

7 The published protocol [20, 21] does not specify how many dilutions were used, so for illustrative purposes, we selected *n*_dilutions_ = 8.

8 We note that real assays may encounter solubility issues with such high compound concentrations, and that the nonideal nature of water:DMSO solutions means that serial dilution of DMSO stocks will not always guarantee all dilutions will readily keep compound soluble. Here, we also presume the DMSO and EDTA control wells are not used in fitting to obtain *p*IC_50_ values.

## References

[1] B. A. Kozikowski, Journal of Biomolecular Screening 8, 210 (2003).

[2] B. A. Kozikowski, Journal of Biomolecular Screening 8, 205 (2003).

[3] X. Cheng, J. Hochlowski, H. Tang, D. Hepp, C. Beckner, S. Kantor, and R. Schmitt, Journal of biomolecular screening 8, 292 (2003).

[4] T. J. Waybright, J. R. Britt, and T. G. McCloud, Journal of Biomolecular Screening 14, 708 (2009).

[5] D. Harris, J. Olechno, S. Datwani, and R. Ellson, Journal of Biomolecular Screening 15, 86 (2010).

[6] R. J. Grant, K. Roberts, C. Pointon, C. Hodgson, L. Womersley, D. C. Jones, and E. Tang, Journal of Biomolecular Screening 14, 452 (2009).

[7] A. Tjernberg, Journalof BiomolecularScreening 11, 131 (2005).

[8] A. Simeonov, A. Jadhav, C. J. Thomas, Y. Wang, R. Huang, N. T. Southall, P. Shinn, J. Smith, C. P. Austin, D. S. Auld, and J. Inglese, Journal of Medicinal Chemistry 51, 2363 (2008).

[9] J. B. Baell and G. A. Holloway, Journal of Medicinal Chemistry 53, 2719 (2010).

[10] L. Di and E. H. Kerns, Drug Discovery Today 11, 446 (2006).

[11] S. L. McGovern, E. Caselli, N. Grigorieff, and B. K. Shoichet, Journal of Medicinal Chemistry 45, 1712 (2002).

[12] S. L. McGovern and B. K. Shoichet, Journal of Medicinal Chemistry 46, 1478 (2003).

[13] B. Y. Feng, A. Shelat, T. N. Doman, R. K. Guy, and B. K. Shoichet, Nature Chemical Biology 1, 146 (2005).

[14] B. Y. Feng and B. K. Shoichet, Journal of Medicinal Chemistry 49, 2151 (2006).

[15] M. Busch, H. B. Thorma, and I. Kober, Journal of Biomolecular Screening 18, 744 (2015).

[16] J. D. Chodera and D. L. Mobley, Annual Review of Biophysics 42, 121 (2013).

[17] C. Kramer, T. Kalliokoski, P. Gedeck, and A. Vulpetti, Journal of Medicinal Chemistry 55, 5165 (2012).

[18] T. Kalliokoski, C. Kramer, A. Vulpetti, and P. Gedeck, PLoS ONE 8, e61007 (2013).

[19] S. Ekins, J. Olechno, and A. J. Williams, PLoS ONE 8, e62325 (2013).

[20] B. C. Barlaam and R. Ducray, Novel pyrimidine derivatives 965, 2009, uS20090054428 A1.

[21] B. C. Barlaam, R. Ducray, and J. G. Kettle, Pyrimidine derivatives for inhibiting Eph receptors, 2010, uS7718653 B2.

[22] G. Xia, S. R. Kumar, R. Masood, S. Zhu, R. Reddy, V. Krasnoperov, D. I. Quinn, S. M. Henshall, R. L. Sutherland, J. K. Pinski, and others, Cancer research 65, 4623 (2005).

[23] C. Bardelle, D. Cross, S. Davenport, J. G. Kettle, E. J. Ko, A. G. Leach, A. Mortlock, J. Read, N. J. Roberts, P. Robins, and E. J. Williams, Bioorganic & Medicinal Chemistry Letters 18, 2776 (2008).

[24] D. Lowe, Drug Assay Numbers, All Over the Place. In the Pipeline:, 2015, http://blogs.sciencemag.org/pipeline/archives/2013/05/03/drug_assay_nL

[25] D. Evanko, Serial dilution woes, 2013, http://blogs.nature.com/methagora/2013/05/serial-dilution-woes.html.

[26] S. Ekins, What it took to get the paper out, 2013, http://www.collabchem.com/2013/05/03/what-it-took-to-get-the-paper-out/.

[27] J. Palmgren, J. Monkkonen, T. Korjamo, A. Hassinen, and S. Auriola, European Journal of Pharmaceutics and Biopharmaceutics 64, 369 (2006).

[28] J. R. Taylor, An Introduction to Error Analysis: The Study of Uncertainties in Physical Measurements (University Science Books, ADDRESS, 1997).

[29] C, Shalizi, Simple Simulation Methods for Quantifying Uncertainty.

[30] L, Walling, N, Carramanzana, C, Schulz, T, Romig, and M, Johnson, ASSAY and Drug Development Technologies 5, 265 (2007).

[31] S, Weiss, G, John, I, Klimant, and E, Heinzle, Biotechnology Progress 18, 821 (2002).

[32] E, Mitre, M, Schulze, G. A. Cumme, F, Rossler, T, Rausch, and H, Rhode, Journal of Biomolecular Screening 12, 361 (2007).

[33] Tecan Genesis Operating Manual, 2001.

[34] Echo 5XX Specifications, 2011.

[35] H, Dong, Z, Ouyang, J, Liu, and M, Jemal, Journal of the Association for Laboratory Automation 11, 60 (2006).

[36] H. Gu and Y. Deng, Journal of the Association for Laboratory Automation 12, 355 (2007).

[37] G. Jones, Journal of Computer-Aided Molecular Design 29, 1 (2015).

[38] J. Wingfield, Impact of acoustic dispensing on data quality in HTS and hit confirmation., 2012.

[39] J. Olechno, S. Ekins, A. J. Williams, and M. Fischer-Colbrie, Direct Improvement with Direct Dilution, 2013, http://americanlaboratory.com/914-Application-Notes/142860-Direct-Improvement-With-Direct-Dilution/.

[40] J. Olechno, S. Ekins, and A. J. Williams, Sound Dilutions, 2013, https://theanalyticalscientist.com/issues/0713/sound-dilutions/.

[41] J. Olechno, J. Shieh, and R. Ellson, Journal of the Association for Laboratory Automation 11, 240 (2006).

[42] R. E. Jones, W. Zheng, J. C. McKew, and C. Z. Chen, Journal of laboratory automation 2211068213491094 (2013).

